# A Kinematic Index for Estimation of Metabolic Rate Reduction in Running with *I-RUN*

**DOI:** 10.1101/2020.08.30.274365

**Authors:** Hamidreza Aftabi, Rezvan Nasiri, Majid Nili Ahmadabadi

## Abstract

In this paper, we target multiple goals related to our passive running assistive device, called *I-RUN*. The major goals are: (1) finding the main reason behind individual differences in benefiting from our assistive device at the muscles level, (2) devising a simple measure for on-line *I-RUN* stiffness tuning, and creating a lab-free simple kinematic measure for (3) estimating metabolic rate reduction as well as (4) training subjects to maximize their benefit from *I-RUN*. Our approach is using some extensive data-driven OpenSim simulation results employing a generic lower limb model with 92-muscles and 29-DOF.

It is observed that there is a significant relation between the hip joints kinematic and changes in the metabolic rate in the presence of *I-RUN*. Accordingly, a simple kinematic index is devised to estimate metabolic rate reduction. This index not only explains individual differences in metabolic rate reduction but also provides a quantitative measure for training subjects to maximize their benefits from *I-RUN*.

The simulation results also re-confirm our hypothesis that “reducing the forces of two antagonistic mono-articular muscles is sufficient for involved muscles’ total effort reduction”. Consequently, we introduce a two-muscles EMG-based metric for the on-line tuning of *I-RUN*.

## Introduction

Passive exoskeletons for metabolic rate reduction ^1^ in running and walking are popular in the research community because of their innovative design as well as the perspective of usability, low weight, and affordable cost^1,3–6^. Nevertheless, as for the active ones, passive exoskeletons have not fulfilled the desired expectations yet. The main reason is lack of a comprehensive model for accurate user-exoskeleton interaction; e.g., predictive forward dynamic models^7–9^. That chiefly stems from high redundancy in our musculoskeletal system^10–12^ and significant individual differences^13, 14^, as well as the complexity of our neural control system^15, 16^. As a result, there is no specific and straight forward method for an exoskeleton design, estimation of its efficacy for a potential user, and on-line adaptation to individuals. Therefore, extensive experiments, laboratory measurements, hand-tunings, and user training are required to benefit from an exoskeleton.

The existing unpowered assistive devices are equipped with elastic elements to save energy in a certain phase of motion and recycle it during another phase. The assistive torque applied by an elastic element is a function of joints’ angles. Therefore, the performance of these devices in metabolic rate reduction and comfort increment is very sensitive to users’ kinematic and the compliance profiles of elastic parts. As a result, **(1)** adaptation of compliant elements (devising a semi-passive exoskeleton)^17, 18^ as well as **(2)** modification of subjects’ kinematic by training^3^ are two solutions for improving the performance of a full-passive exoskeleton. To materialize these two approaches for an assistive device, a low-cost and simple compliance adaptation method and a quantitative straight-forward kinematic measure for subject training are required. Such a measure can be potentially employed to estimate exoskeletons’ metabolic rate reduction for an individual as well; it is a substitute for laboratory-bounded costly devices like gas analyzers. In this paper, we target these requirements for our previously designed exoskeleton device named *I-RUN*^1^.

Previously, we reported the first unpowered exoskeleton for metabolic rate reduction during running; on average 8 ± 1.5% at 2.5*m/s*^1^. However, we observed high individual differences in metabolic rate reduction, and there was no clear explanation for that. The only thing we knew was that there is a certain relation between the metabolic rate reduction, the compliance coefficient, and the subjects’ kinematic during running. To investigate and exploit that relation for the above-mentioned purposes, we employ some extensive detailed simulations.

Using simulations for exoskeleton design and analysis, as a partial substitute for costly experiments, has gained momentum in recent years^19–23^. For instance, based on simulation results, it is suggested that a shifted exoskeleton torque profile for the hip joint is very effective for reducing the metabolic cost of running^21^; based on that an active exosuit device was devised24. As another example, Jackson *et al.*^23^ explained the exoskeleton effects on the biomechanical parameters by a forward simulation approach.

Here, the same strategy is taken and by using a simulation-based approach, we investigate the relation between metabolic rate reduction and other contributors; namely elastic element stiffness, exoskeleton torque profile, biological moment, muscles’ forces, and gait kinematic. Accordingly, a simple kinematic index for subject training and estimation of metabolic rate reduction is devised. This index explains individual differences in metabolic rate reduction and provides a quantitative measure for subjects to maximize their benefits from *I-RUN*. Besides, we study if there is a relation between the metabolic rate and all muscles’ effort. Computation of all muscles’ effort requires each muscle’s electromyography (EMG) signal, that makes it inapplicable in practice. To resolve this problem, inspired by Nasiri *et al.*^25^, we test our hypothesis that “the effort of two antagonistic mono-articular muscles at the target joint has a linear correlation with all muscles’ effort”. Having this hypothesis validated, it is shown that, reducing the EMG signal of only two antagonistic mono-articular muscles is sufficient for whole muscles’ effort reduction. In addition, it is demonstrated that the compliance value resulting in the minimum effort of two antagonistic mono-articular muscles is a good sub-optimal for the metabolic rate reduction. These findings simplify online exoskeleton adaptation.

## Methods & Materials

### Experimental data

Dataset of 10 subjects reported by Hamner *et al.*^26^, was used in OpenSim^27^ to run a muscle driven simulation. Due to technical issues, we were only able to use the data of 7 subjects in OpenSim 4.0. As mentioned by Hamner *et al.*^26^, subjects have an average age, height, and mass of 29 ± 5*y*, 177 ± 4*cm*, and 70.9 ± 7.0*kg*, respectively. This dataset contains marker positions and ground reaction forces (GRF) of running gait at four different speeds: 2.0, 3.0, 4.0, and 5.0*m/s*. Although the experimental validation of *I-RUN*^1^ was at 2.5*m/s*, here, due to limitations of the mentioned dataset, we had to choose either 2.0*m/s* or 3.0*m/s*. As it is reported by Schache *et al.*^28^, for speeds higher than 2.5*m/s*, the posture, dynamics, and trajectory of the joints are significantly changed; hence, 2*m/s* was chosen for our simulation study.

### Dynamic Simulations

We used a 12 segment musculoskeletal model with 29 DOF^29^ in OpenSim to run a musculoskeletal simulation; this model covers frontal, lateral, and sagittal movements. Lower extremity and back joints of this model were actuated by 92 hill-type muscle-tendon units^30^, and arms were controlled by ideal actuators. The generic model was scaled to create a model consistent with each subject’s anthropometry. Scaling was done by minimizing the difference between experimental markers^2^ and their corresponding virtual markers^3^ in a static pose. To calculate each joint angle, we used an inverse kinematics procedure. Unlike the scaling, which uses the marker trajectories of the static pose, the inverse kinematics algorithm uses the marker trajectories of the running gait as an input to minimize the difference between experimental and virtual markers.

*I-RUN* applies a torque on hip joints proportional to the difference between left and right hip angles. Accordingly, this mechanism couples two hips with each other using a torsional spring; i.e., BLS in Fig. 1. To model this assistive device, two ideal coordinate actuators were added to the hip joints. For evaluating our exoskeleton in different stiffnesses of BLS, we defined a factor named spring coefficient (*α*); *α* scales the inverse dynamic stiffness (*K_inv_*). *K_inv_* is the stiffness value for the exoskeleton device that minimizes 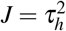 where *τ_h_* is the biological hip moment. In this case, the assistive torque of the exoskeleton is written as *τ_a_* = *αK_inv_*Δ*θ* where *τ_a_* represents the assistive torque and Δ*θ* is the difference between left and right hip angles. It is obvious that *α* = 0 and *α* = 1 refer to no assistive torque and full possible average hip moment compensation conditions, respectively.

**Figure 1.**
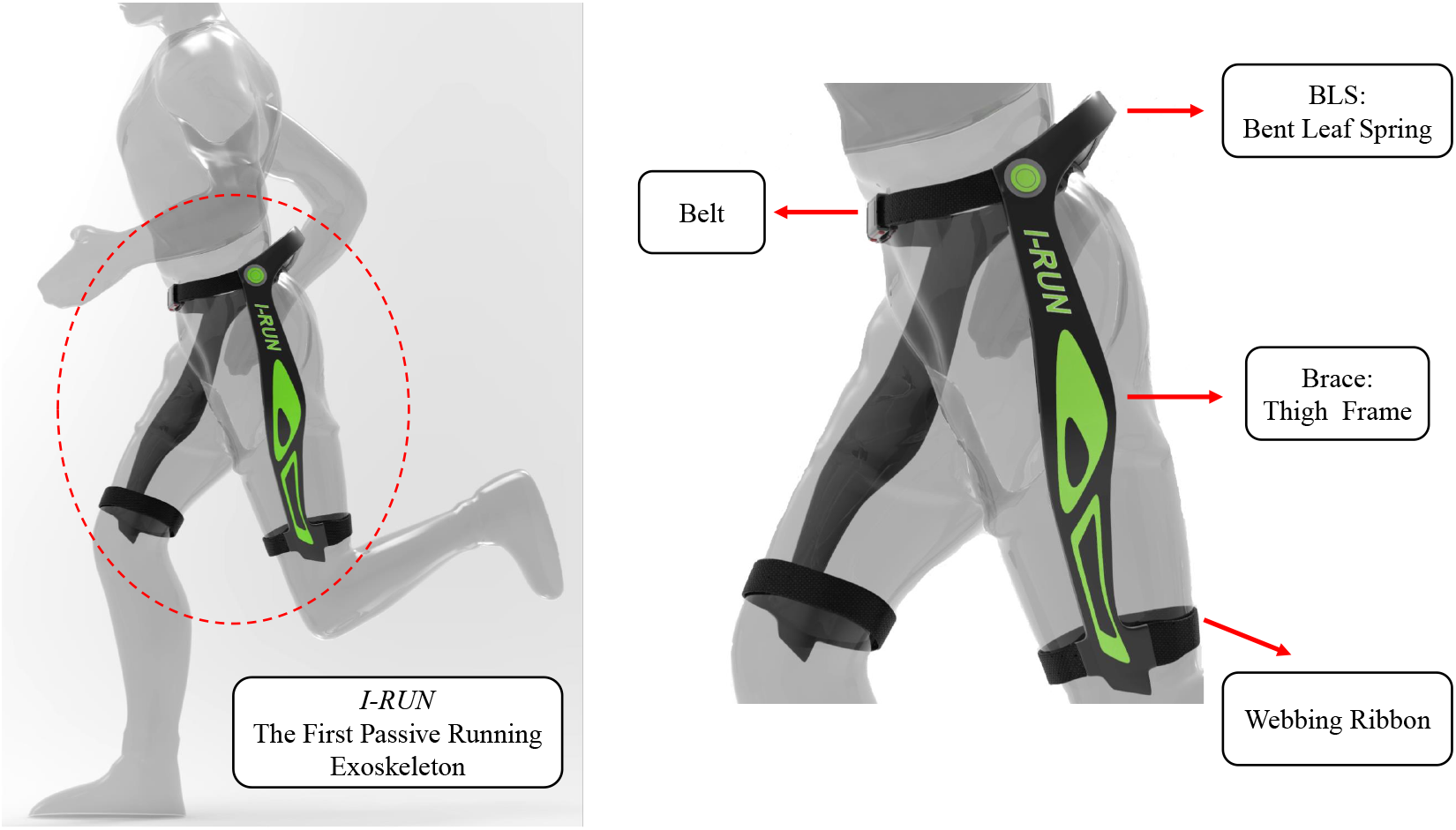
Industrial design of *I-RUN*^1, 2^. *I-RUN* resulted in on average 8% metabolic rate reduction in ten healthy active subjects at 2.5*m/s* running speed.

We used the residual reduction algorithm (RRA)^27^ to calculate the moment of each joint. RRA algorithm modifies the torso mass center of the subject’s specific model. Besides, it allows for minor changes in joint angles (less than 2*deg*), previously calculated from the inverse kinematics, to create a model that is more compatible with the ground reaction forces dataset. We employed the computed muscle control algorithm (CMC)^31^ to calculate the muscles’ forces. CMC minimizes the sum of the square of muscles’ forces while driving the musculoskeletal model toward desired kinematics. CMC objective function can be written as^32^:

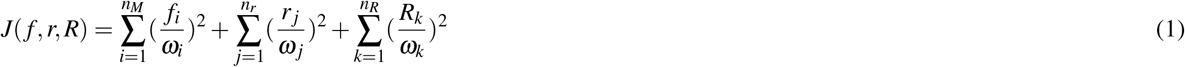

where *f*, *r*, and *R* are muscle forces, reserve torques, and residual torques, respectively; *n_M_* = 92, *n_r_* = 29, and *n_R_* = 6. Besides, *ω_i,j,k_* refer to their corresponding weights. Reserve actuators (*r*) are responsible for the model’s strength by applying small torques in joints, while residual actuators model 6 degree of freedom (3 translational, 3 rotational) between pelvis of the generic model and the ground. Weights of residual and reserve actuators are chosen small enough to ensure a high cost in the objective function.

### Data analysis

For each subject, the metabolic cost at each spring coefficient was calculated using the method presented by Uchida *et al.*^33^. The total metabolic cost was computed by integrating the instantaneous metabolic cost over one gait cycle and divided by the stride period. Besides, to normalize the results, the total metabolic cost was divided by each subject’s weight.

Average muscle force and moment were calculated by integrating the force and absolute moment of the time series data and divided by the stride period and the subject’s mass.

In this paper, normalized muscle force is defined as the muscle force divided by its maximum isometric force. In addition, the instantaneous all muscles’ effort is defined as the summation of the square of all normalized muscles’ forces^34, 35^. And, the instantaneous antagonistic mono-articular muscles’ effort at the hip joint is the summation of the square of the normalized Psoas and Gluteus Maximus muscles’ forces. Finally, muscles’ effort was computed by integrating the instantaneous muscles’ effort over one gait cycle and divided by the stride period.

### Statistics

At each spring coefficient, means and s.e.ms of net metabolic rate, average joint moment, average muscle force and, muscle effort were calculated across the subjects. Due to previous observations that the metabolic rate is a nonlinear function of different stiffnesses^4, 36^ and the hypothesis that the muscles’ effort is a quadratic function of muscles’ forces^34, 35^, we performed a mixed model, three-factor ANOVA (random effect: participant; fixed effects: spring coefficient and square of spring coefficient; significance level = 0.05; JMP Pro) to evaluate the effect of the spring coefficients on the metabolic rate and muscles’ effort. We also carried out a two-factor ANOVA test (random effect: participant; fixed effects: spring coefficient; significance level = 0.05; JMP Pro) to assess the effect of the spring coefficients on the joint moment and muscle force. Besides, a least-square method was employed to (1) fit a second-order polynomial to the mean data of the metabolic rate and muscles’ effort and (2) the best linear relation between potential kinematic indexes and the metabolic rate reduction in the optimum stiffness.

For post hoc analysis, we utilized a paired two-sided t-test for multiple comparisons with Holm-Bonferroni correction (significance level = 0.05; Scikit-Learn, Python 3.6) in order to compare the spring coefficients with each other and find out which spring coefficient yields a significant change in the net metabolic rate and muscle effort. Finally, a Shapiro-Wilk test (significance level = 0.05; JMP Pro) was employed to ensure the appropriateness of the tests in which normality is a necessary condition.

### Results evaluation

Finally, to assess the validity of our results, we evaluated different parameters, such as peak residual moments, peak residual forces, and peak reserve actuator torques. In all conditions and for all subjects, maximum residual moments(forces) were less than 40*Nm*(12.5*N*); the maximum residual force was much less than 5% of the peak magnitude of experimental GRF^37^. And, the peak of reserve actuator torques were less than 20*Nm*. Besides, by comparing the kinematics of simulations and experimental data, it is observed that the maximum RMS deviation is lower than 2*deg*. As it is mentioned before, the maker locations on the model were scaled to fit with the real dataset; RMS markers error did not exceed 2*cm*. All of these values were less than OpenSim best practice thresholds^38^, which approves the preciseness of our simulations.

### Assumptions, limitations, and considerations

**Weight**: The mass of the assistive device is ignored in this study. The mass of *I-RUN* is almost 1.8 kg^1^, which is negligible compared to the average weight of the subjects (77 ± 7.0*kg*). **Solver:** The main weakness of static optimization approaches is that they assume that the subject’s motions are almost fixed before and after adding/wearing an exoskeleton device. Hence, the results of this method are reliable for the cases that the joint trajectories are not significantly changed by exoskeleton augmentation. Besides, the static optimization methods suffer from the lack of smoothness in their extracted torques/forces profiles. Accordingly, the variation among different types of static optimization approaches comes from their data-driven solutions for resolving the biomechanical variables, and their chosen cost function or redundancy resolution method. Among all of the static optimization toolboxes for neuromuscular model resolution, computed muscle control (CMC)^31^ provides smoother torques/forces profiles. And, due to the promising and reliable results of CMC, it has gained much attention in recent years for analyzing the biomechanical effects of adding exoskeleton devices. **Other limitations**: for the sake of simplicity and without losing generality, we did not model muscle fatigue, and co-activation; assuming that the co-activations don’t change with and without *I-RUN* because of keeping the gate the same.

## Results & Discussion

### Metabolic rate reduction

To find a metric for the metabolic rate reduction, as the first step, we compute the optimum stiffness for the metabolic rate reduction in our model. Accordingly, we plot the average metabolic cost of seven subjects (in Fig. 2) versus different spring coefficients; from *α* = 0 to *α* = 1.5 with the step size of 0.1. As can be seen in Fig. 2, the spring coefficient value that minimizes the metabolic rate is at *α\** = 0.6; the mean metabolic rate reduction at *α\** = 0.6 is 4.62%. It is worth noting that *α*\* = 0.6 (calculated by exhaustive search) is close to the spring coefficient which was experimentally reached by try-and-error^1^. Interestingly, similar to the research by Collins *et al.*^4^, the metabolic cost versus the spring coefficient in Fig. 2 is a quadratic function.

**Figure 2.**
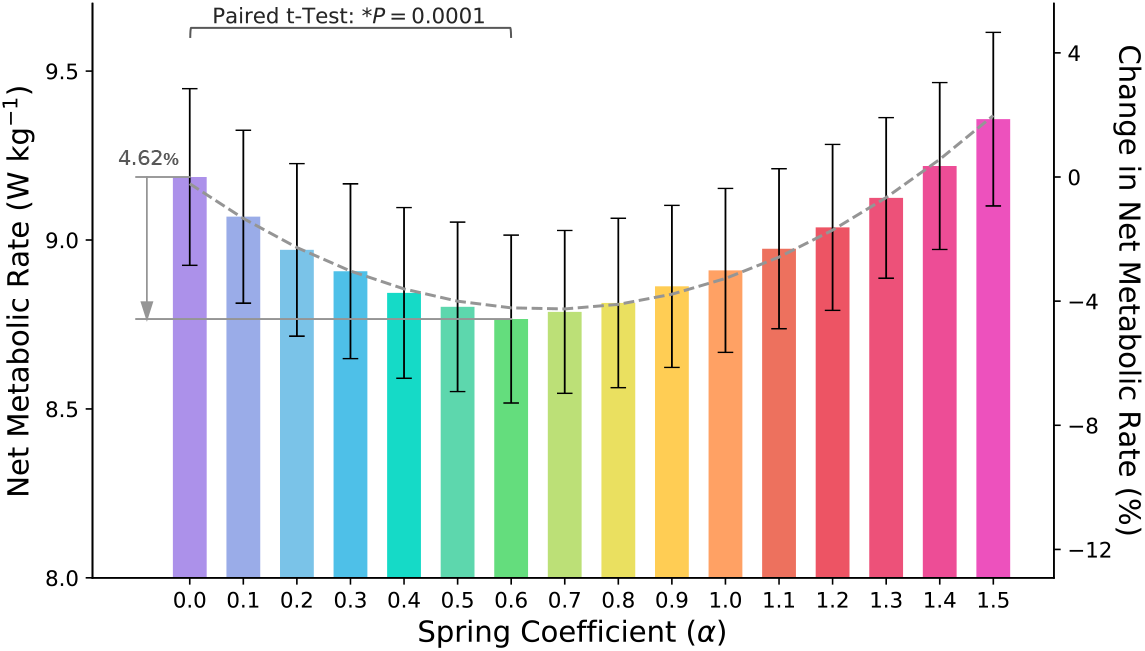
Metabolic rate. Metabolic rate versus the stiffness coefficient (N = 7; ANOVA with second order model; random effect: participant; fixed effects: spring coefficient and square of spring coefficient; *P_scoef_* < 0.0001, *P*_*scoef*^2^_ < 0.0001). The maximum metabolic rate reduction is 4.62 ± 0.29% (mean s.e.m) at *α*\* = 0.6 (paired two-sided t-test with correction for multiple comparisons; *P* = 0.0001). The dashed line is the quadratic best fitted profile to average metabolic costs (*R*^2^ = 0.98, *P* < 0.0001).

### Biological joints’ moments

Here, we intend to analyze the relation between metabolic rate reduction and average biological moment reduction. Accordingly, we plot the biological moment of all joints (hip, knee, and ankle) versus the gait cycle at different spring coefficients in Fig. 3. As can be seen, only the hip torque profile is affected by exoskeleton augmentation, and the torque profiles of the other joints (knee and ankle) remain almost fixed. This is an obvious observation since the kinematic of the gait is assumed to be a fixed function of time with/without exoskeleton torque augmentation at the hip joint; see the joint kinematics in Fig. 10.

**Figure 3.**
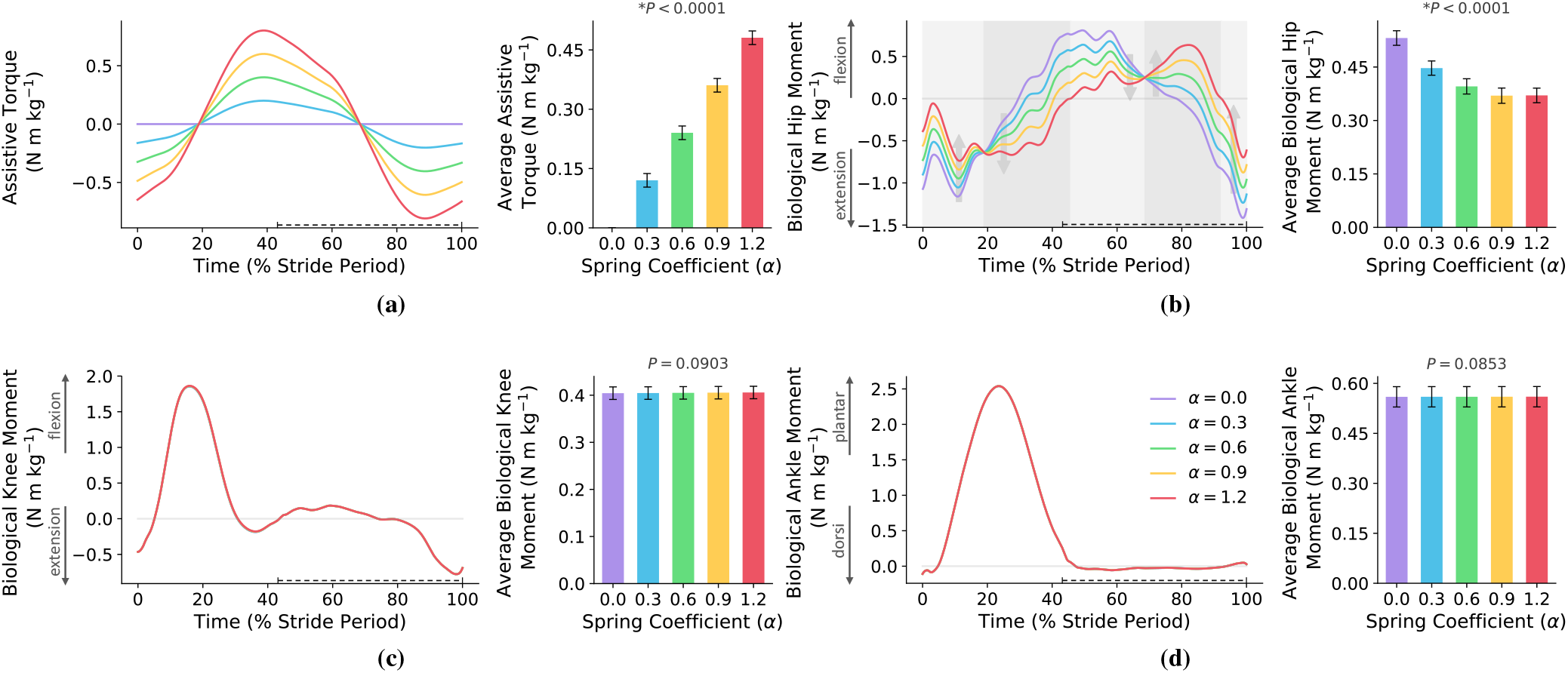
Joints’ torques. This figure shows the effects of exoskeleton augmentation on the joints’ biological moments and their average over the whole stride. The start of the gait cycle is at the heel-strike instant of the right leg and all of the figures are for the right leg. The swing phase is shown with a dashed line over the horizontal axes. (a) is the exoskeleton torque profile for different stiffness coefficients. (b-d) show the biological hip, knee, and ankle moments. As it is clear, the exoskeleton augmentation only affects the biological hip moment and the moments of the knee and ankle are not changed. The background color code for sub-figure (b) is: light and dark gray mean biological hip moment improvement and disruption/increment, respectively. Moreover, the minimum of the average biological hip moment is on *α* = 1 where it has 31% reduction compared to no exoskeleton case. And, the reduction of the average biological hip moment for *α\** = 0.6 is 26%. N = 7; bars, mean; error bars, s.e.m; P-values, ANOVA (random effect: participant; fixed effects: spring coefficient; significance level = 0.05).

Based on Fig. 3, compared to the no exoskeleton case (*α* = 0), the exoskeleton augmenting (*α >* 0) disturbs or improves the biological hip moment in the dark and light gray regions, respectively. In the dark gray regions, exoskeleton torque augmentation either increases the amplitude of biological hip moment or reverses its sign. Increasing the hip biological moment increases the muscles’ effort. And, any variation in the sign of the biological moment changes the muscles’ activation patterns^4^. In light gray regions, the exoskeleton torque augmentation (*α >* 0) reduces the biological hip moment while its sign remains the same, which is the best possible condition for improving the muscles’ effort without any changes in the muscles’ activation patterns. Although dark gray and light gray regions are the same for all *α >* 0, for a certain spring coefficient (*α* = 1) we have on average the most biological hip moment reduction; *α* = 1 is the solution for inverse dynamics torque minimization.

Based on this observation, the spring coefficient (*α\** = 0.6) that minimizes the metabolic rate is much lower than the one which minimizes the biological hip moment (*α* = 1). In fact, this simulation supports the hypothesis that “a moderate stiffness is sufficient for metabolic rate reduction”^4^. In other words, the stiffness coefficient that minimizes the metabolic rate is far lower than the value that minimizes the average biological hip moment. The reason for this difference relies on the overall scope of the average biological hip moment index. Reducing the average biological moment does not care about the biomechanics of the body. It is due to the existence of bi-articular muscles, antagonistic behavior, and high redundancy in the number of muscles. Therefore, in the next subsection, we check if minimizing individual muscles’ forces results in a better estimation of the optimum spring coefficient of *I-RUN*.

### Individual muscles’ forces

In this subsection, we study if there is a correlation between the metabolic rate reduction and individual muscles’ forces trajectory and their average over one gait cycle. To do so, the force profiles of nine main muscle branches in the presence of exoskeleton augmentation are illustrated in Fig. 4 and Fig. 5. Fig. 4 shows the four main muscles contributing at the hip joint (Psoas in Fig. 4a, Gluteus Maximus in Fig. 4b, Rectus Femoris in Fig. 4c, and Semimembranosus in Fig. 4d) and the other five muscles are plotted in Fig. 5 (Vastus Laterialis in Fig. 5a, Bicep Femoris in Fig. 5b, Gastrocnemius Medialis in Fig. 5c, Tibialis Anterior in Fig. 5d, and Soleus in Fig. 5e). The exact placement of the nine main muscle branches are shown in Fig. 5f.

**Figure 4.**
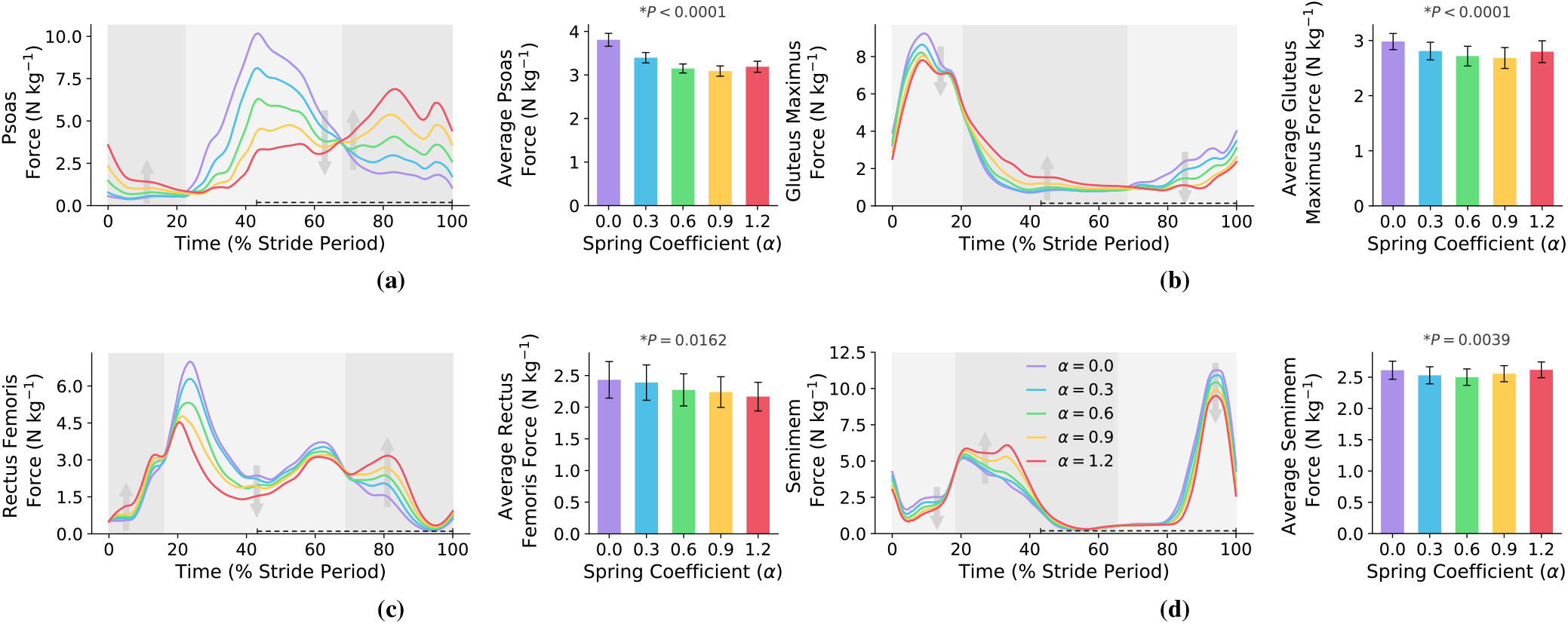
Hip muscles’ forces. This figure shows the effects of exoskeleton augmentation on the main hip muscles’ forces and their average over the whole stride. The start of the gait cycle is at the heel-strike instant of the right leg and all of the figures are for the right leg. The swing phase is shown with a dashed line over the horizontal axes. The background color code for all of the sub-figures is: light(dark) gray means muscle force reduction(increment). As can be seen, exoskeleton augmentation changes the force profiles of all hip muscles. (a,b) show main hip antagonistic mono-articular muscles where Psoas is a hip flexor and Gluteus Maximus is a hip extensor. The minimum of average Psoas and Gluteus Maximus forces are on *α* = 0.8 which yields to 19% and 10% reductions. And, the reductions of average Psoas and Gluteus Maximus forces for *α\** = 0.6 are 17.4% and 8.9%, respectively. (c,d) show main hip antagonistic bi-articular muscles where Rectus Femoris and Semimem are hip flexor and extensor, respectively. The minimum of average Rectus Femoris and Semimem forces are on *α* = 1.4 and *α* = 0.6 which yields to 11.7% and 4.6% reductions. And, the reductions of average Rectus Femoris and Semimem forces for *α\** = 0.6 are 6.5% and 4.2%, respectively. N = 7; bars, mean; error bars, s.e.m; P-values, ANOVA (random effect: participant; fixed effects: spring coefficient; significance level = 0.05).

**Figure 5.**
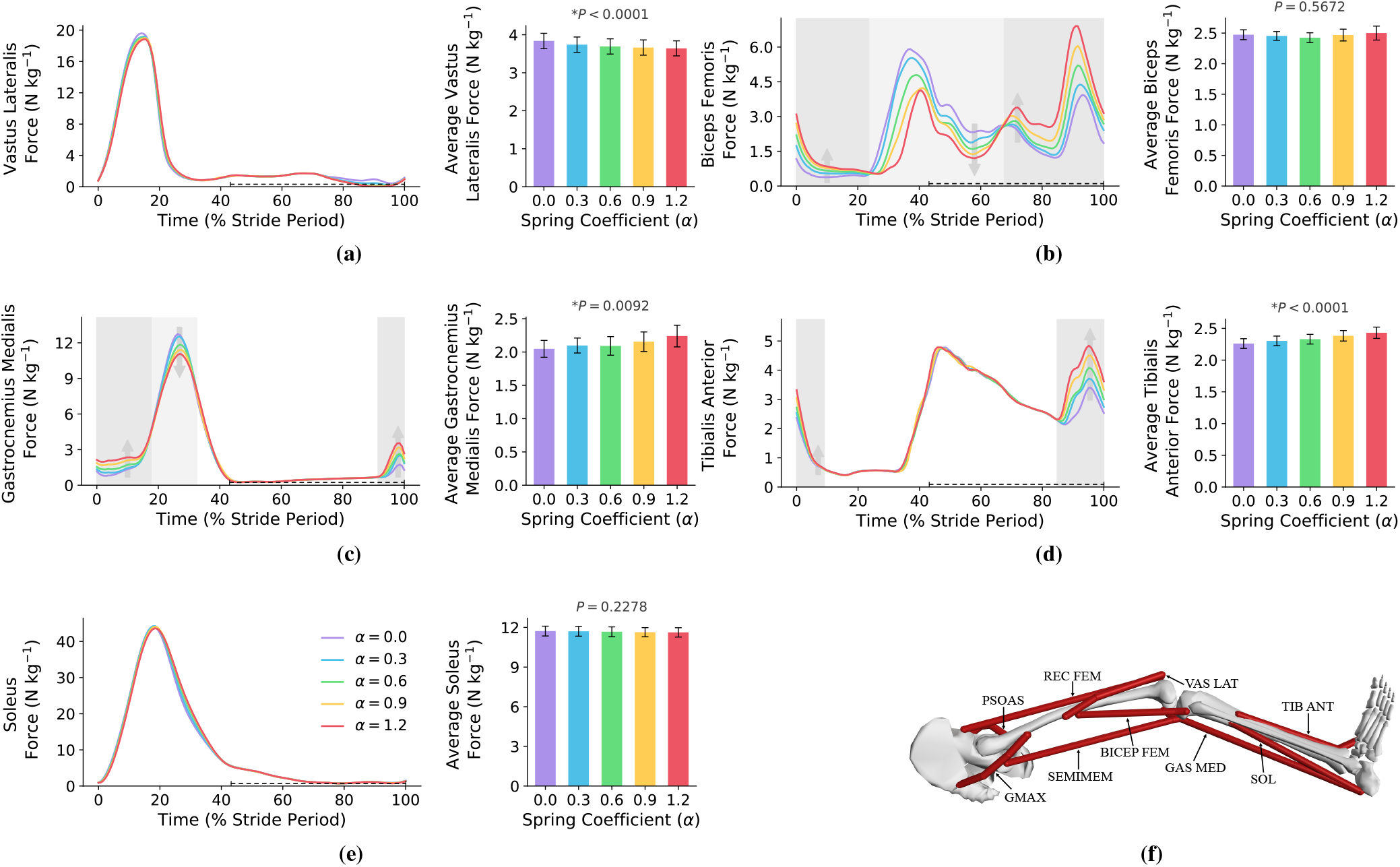
Other muscles’ forces. (a-e) sub-figures show the effects of exoskeleton augmentation on the main muscles’ forces and their average over the whole stride except for the main hip muscles. The start of the gait cycle is at the heel-strike instant of the right leg and all of the figures are for the right leg. The swing phase is shown with a dashed line over the horizontal axes. The background color code for all of the sub-figures is: white means no special effect, light gray means muscle force reduction, and dark gray means muscle force increment. As can be seen, the force of Soleus and Vastus Laterialis have no significant changes by exoskeleton augmentation. Besides, despite the significant changes in the Bicep Femoris force profile, the average force of this muscle shows no significant change in different spring coefficients; minimum on *α* = 0.6 with 2% reduction. In addition, the optimum spring coefficient for Gastrocnemius Medialis and Tibialis Anterior are on *α* = 0. And, the minimum average force for Vastus Lateralis and Soleus are on *α* = 1.5 with 5.6% and 1.2% reductions compared to no exoskeleton case. Moreover, the average forces of these muscles have 3.8% and 0.5% reductions on *α\** = 0.6, respectively. N = 7; bars, mean; error bars, s.e.m; P-values, ANOVA (random effect: participant; fixed effects: spring coefficient; significance level = 0.05). (f) shows the placement of the nine main branches of the muscles in the lower limb.

For hip muscles (Fig. 4), the exoskeleton augmentation leads to flexor muscles’ (Psoas and Rectus Femoris) forces increment approximately between 0% – 20% and 70% – 100% of the gait cycle while the extensor muscles’ (Gluteus Maximus and Semimembranosus) forces are reduced in these phases. The reverse behavior can be seen for 20% – 70% of the gait cycle where the hip flexor(extensor) muscles’ forces are reduced(increased). This behavior can be explained by comparing the exoskeleton torque profile in Fig. 3a with the hip muscles’ forces in Fig. 4; there is an interesting correlation between these two figures. Accordingly, exoskeleton flexion(extension) torque decreases the flexor(extensor) muscles’ forces while amplifies the extensor(flexor) pair antagonistic muscle at the same time. Analyzing the average force over one gait cycle w.r.t. the metabolic rate diagram does not provide a logical correlation for the metabolic rate reduction.

Fig. 5 also shows the force of other muscles; those muscles which are not connected to the hip joint. Based on Fig. 5, except Bicep Femoris muscle, exoskeleton augmentation leads to insignificant changes in the force of the other muscles such that the forces of Vastus Lateralis and Soleus are almost fixed. Although exoskeleton augmentation does not affect the biological moment profile of knee and ankle joints (see Fig. 3), it minimally changes the muscles’ forces that are not connected to the hip joint. This is due to the function of bi-articular muscles in the lower limb joints; torque distribution over the whole muscles. Accordingly, any variation in a single joint biological moment changes the force profiles of whole lower limb muscles; the changes are reduced by getting away from the augmented joint.

Since the metabolic rate reduction has some relations with individual muscles’ average forces reduction (e.g., Semimem), a force-based index may potentially provide us with a measure for the metabolic rate reduction. In the next subsection, we study the relation between the whole muscles’ effort (which is defined upon the muscles’ forces) and metabolic rate reduction.

### Whole muscles’ effort

The whole muscles’ effort is a unified index for muscles’ forces redundancy resolution, and the static optimization methods in OpenSim utilize this index^32^. Besides, momentum minimization does not consider force re-distribution among the muscles; while muscle effort minimization is a candidate index for redundancy resolution and force distribution among the muscles. Therefore, we check if the whole muscles’ effort index minimization leads to the maximum metabolic rate reduction. However, the problem of computing the whole muscles’ effort in the physical experiments is that it requires the force profiles of all muscles. To resolve this issue, previously, we presented the hypothesis that “reducing the efforts of two antagonistic muscles is sufficient for whole muscles’ effort reduction”^25^. Accordingly, instead of using the efforts of 92 muscles in the lower limb model, we can rely on the effort of two antagonistic mono-articular muscles at the targeted joint. However, this hypothesis was studied on a 2-DOF model with 6 muscles (four mono-articular and two bi-articular). Therefore, first, we should check if our hypothesis is valid in a complex and general model of the human lower limb with 29-DOF and 92 muscles.

The whole muscles’ effort versus spring coefficient is plotted in Fig. 6a. In the hip joint, the main antagonistic mono-articular muscles are Psoas and Gluteus Maximus and the efforts of these muscles are also reported in Fig. 6b. Comparing Fig. 6a and Fig. 6b indicates that the average of “all muscles’ effort” and “two main hip antagonistic mono-articular muscles’ effort” are correlated functions of spring coefficient with the same optimum; the average mismatching error for 16 spring coefficient and all subjects is 0.89 ± 1.96% and the linear correlation criteria is 91%; see Fig. 6 and Fig. 7. These results provide support for the correctness of our proposed hypothesis in a complex and generic model of the human lower limb. Using this hypothesis, one can minimize the whole muscles’ effort by utilizing the force of only two mono-articular antagonistic muscles. Accordingly, due to the monotonic relation between muscles’ forces and root-mean-square (RMS) of the EMG signals^39, 40^, we can drastically reduce the number of EMG sensors required in practice; only two EMG sensors are sufficient for *I-RUN* stiffness adaptation.

**Figure 6.**
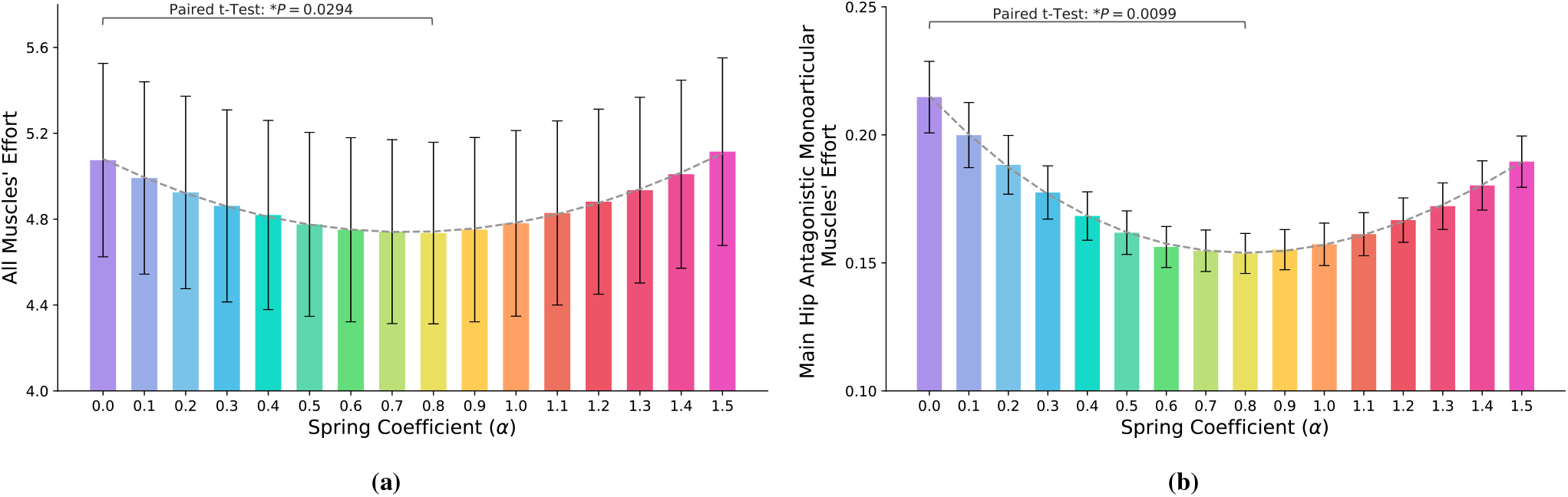
Muscles’ effort. Figure (a) shows all 92 muscles’ effort versus the stiffness coefficient (N = 7; ANOVA; random effect: participant; fixed effects: spring coefficient and square of spring coefficient; *P_scoef_* = 0.0343, *P*_*scoef*^2^_ < 0.0001). The optimum spring coefficient for the whole muscles’ effort reduction is on *α*^#^ = 0.8 (paired two-sided t-test with correction for multiple comparisons; *P* = 0.0294). The dashed line is the quadratic best-fitted profile to average muscles’ effort (*R*^2^ = 0.99, *P* < 0.0001). Figure (b) shows main hip antagonistic mono-articular muscles effort versus the stiffness coefficient (N = 7; ANOVA; random effect: participant; fixed effects: spring coefficient and square of spring coefficient; *P_scoef_* < 0.0001, *P*_*scoef*^2^_ < 0.0001). The optimum spring coefficient for the mono-articular muscles’ effort reduction is on *α*^#^ = 0.8 (paired two-sided t-test with correction for multiple comparisons; *P* = 0.0099). The dashed line is the quadratic best-fitted profile to average muscles’ effort (*R*^2^ = 0.99, *P* < 0.0001). Comparing (a) and (b) shows that stiffness coefficient optimization based on feedback from mono-articular muscles leads to whole muscles’ effort reduction.

**Figure 7.**
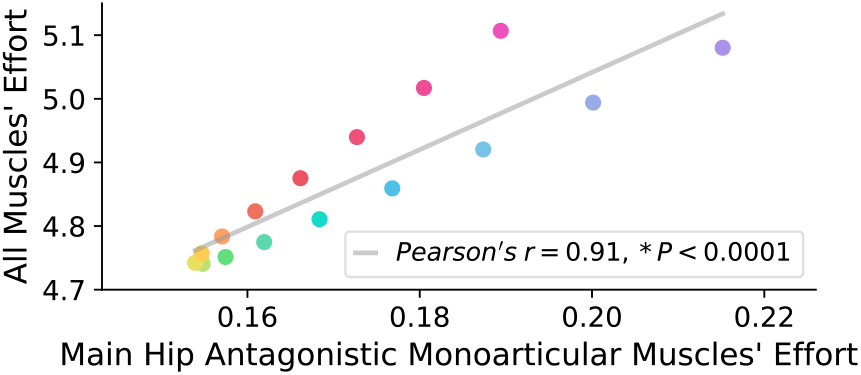
Correlation between all muscles’ effort and two antagonistic mono-articular hip muscles’ effort; the color of each point in this plot is the same with its column color in Fig. 6. The P-value for this linear relation is *P* < 0.0001 and “Pearson’s r-value” for this linear correlation is *r* = 0.91, which shows a certain linear correlation between these two parameters. Interestingly, near to the optimum spring coefficient, the deviations from the linear estimation are minimum and by getting away from the optimum stiffness coefficient the deviations increase.

Comparing Fig. 2 and Fig. 6 shows that the optimum stiffness coefficient suggested by muscles’ effort reduction is different from the value that minimizes the metabolic rate. However, compared to the other candidates as biological hip moment and individual muscles’ effort, it suggests a much closer and reasonable solution for metabolic rate minimization. The suggested spring coefficient (*α*^#^ = 0.8) is a sufficient/reliable sub-optimal for metabolic rate reduction using only two EMG sensors. Compared to the indirect calorimetry, EMG sensors are much easier to use. In addition, they provide us with faster dynamics that not only reduce the experiment time but also significantly improve the speed of adaptation methods compared to gas-analyzer-based approaches; e.g., human-in-the-loop^41, 42^.

### A kinematic index for metabolic rate reduction

*I-RUN* torque is a linear function of the difference between hip angles; i.e., the gait kinematics directly changes the torque profile. Hence, the hip angle and gait kinematic have an undeniable effect on the metabolic rate reduction. Accordingly, a kinematic index has more chance to show the harmony between the gait and the passive exoskeleton torque to maximize the metabolic rate reduction. As it is discussed in **Biological joints’ moments** subsection, a proper torque profile is the one that reduces the amplitude of the biological hip moment without changing its sign. To study the importance of gait kinematics on the exoskeleton torque profile, we compare the biological hip moment, exoskeleton torque, and hip angles of two extreme cases in Fig. 8a and Fig. 8b; i.e. the subjects with the highest and lowest metabolic rate reductions.

**Figure 8.**
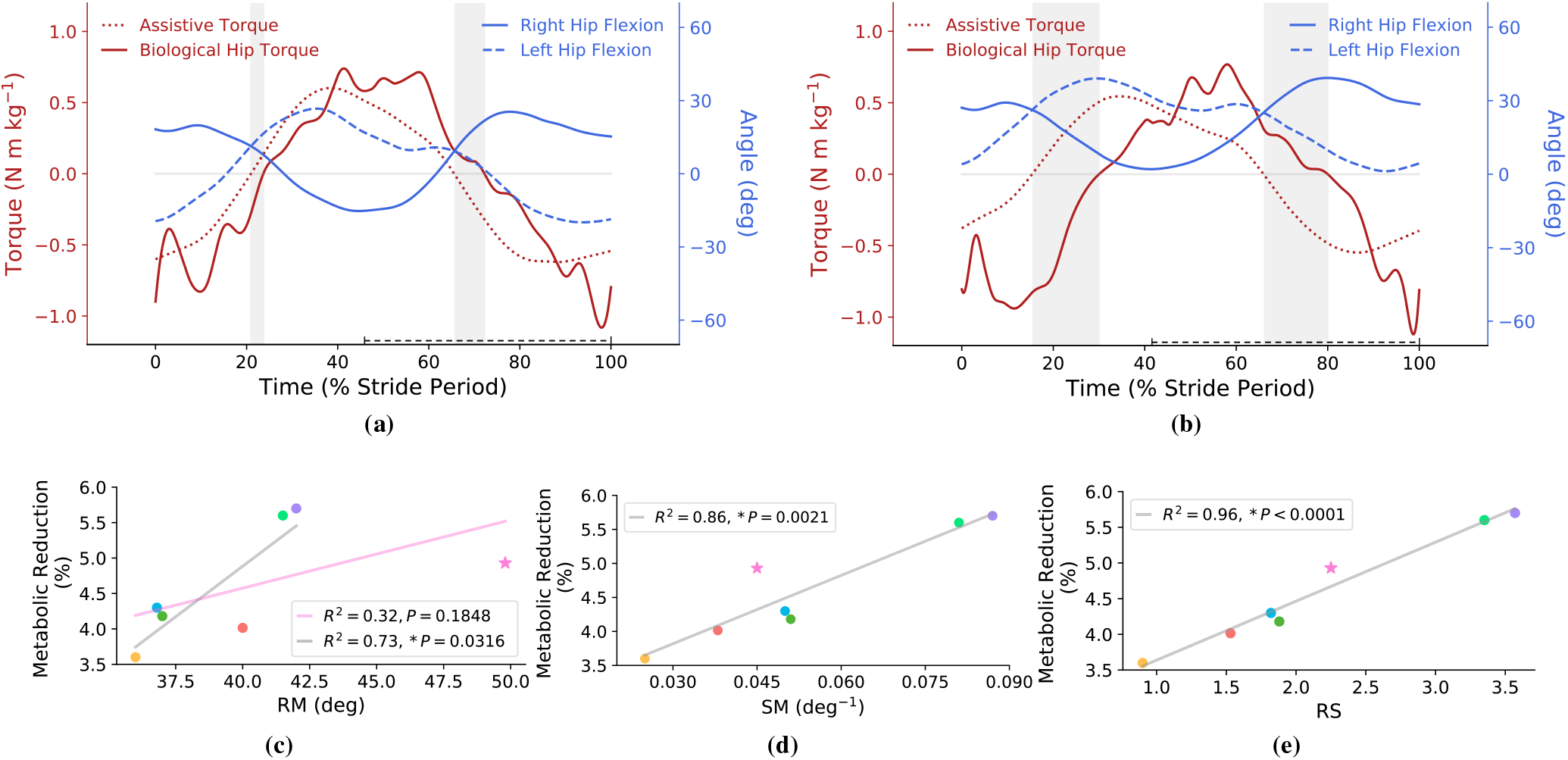
Kinematic analysis. (a) and (b) compare the normalized biological hip torques, hip joints’ angles, and normalized assistive torques for the subjects with the highest and lowest metabolic rate reductions. The start of the gait cycle is at the heel-strike instant of the right leg, and the torque profiles in (a) and (b) are for the right leg. The swing phase is shown with a dashed line over the horizontal axes. Gray areas define the difference between the zero points of the biological hip torque and assistive torque. As it is clear, gray areas are much narrower for the subject with the highest metabolic rate reduction. (c-e) compare three different defined kinematic indexes for estimating the metabolic rate reduction. In figures (c-e), each color indicates one subject. As can be seen, P-value and goodness of fit are improved from RM, SM, and RS = RM × SM, respectively. In (c), the linear regression (pink line) cannot properly estimate the metabolic rate such that there is an outlier (pink star) in this figure; by excluding the outliers based on Jack-knife criterion (significance level = 0:05), the goodness of fit of the linear regression (gray line) is improved. However, in (d) and (e), the outlier approaches to the linear regression and the goodness of fit improves further. Accordingly, the best index for having a good linear relation between metabolic rate reduction and kinematic parameters is RS.

By comparing Fig. 8a and Fig. 8b in the gray regions, it is concluded that if the zero points of exoskeleton torque and the biological hip moment are close to each other, the exoskeleton torque reduces the amplitude of biological hip moment without changing its sign. The zero points of the exoskeleton torque are equivalent to the zero value of the difference of hip angles. Now the question is, how we can change the zero point of exoskeleton torque by kinematic modification?

There are two parameters that contribute to moving the zero point of exoskeleton torque towards the biological hip moment; (1) symmetry of motion (SM) and (2) range of motion (RM) which are depicted in Fig. 10a. The mathematical definitions of SM and RM are 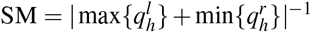 and 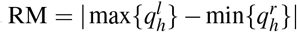 where 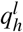 and 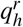 are the angles of the left and the right hip joints. We also define another index (RS) which considers both symmetric motion (SM) and rang of motion (RM) at the same time as RS = RM × SM.

To study the relation between RM, SM, RS, and metabolic rate reduction, we plot the metabolic rate reduction of each individual subject (at the optimum stiffness coefficient *α\** = 0.6) versus RM, SM, and RS in Fig. 8c, Fig. 8d, and Fig. 8e, respectively. In these figures, the metabolic rate reduction is estimated using linear regressions. In Fig. 8c, the stared subject (shown by the pink star) is an outlier for the linear estimation, accordingly, a second linear regression line is also plotted by excluding the outlier subject. However, the stared subject is not an outlier for the SM and RS indexes. Hence, compared to RM, SM and RS are better indexes for estimating metabolic rate reduction. And, among all of the indexes, RS provides the best linear estimation for all of the subjects including the stared one. Therefore, it is concluded that RS^5^ is a rich and representative index for estimating the metabolic rate reduction.

The optimum spring coefficient for most of the subjects is *α\** = 0.6, hence, we can conclude that the optimum spring coefficient is independent of the RS index. It means that changes in the RS index by training for improving metabolic rate reduction shall not change the optimum stiffness.

To study the generality of the RS index, it is required to check if this index is also applicable for the spring coefficients rather than the optimum one. Accordingly, we plot the R-square and the slope of RS linear fitting v.s. different spring coefficients in Fig. 9a and Fig. 9b, receptively. Fig. 9a shows that metabolic rate reduction can be linearly estimated by RS index between *α* = 0.3 and *α* = 1.1. Interestingly, the highest *R*^2^ is at *α\** = 0.6 and for the values around it, the linear fitting is sufficient. Fig. 9b also shows that in all of the spring coefficients the slope of the linear relation is positive; i.e., improvement in RS increases the metabolic rate reduction. In addition, it is concluded that by increasing the spring coefficient, variation of metabolic rate w.r.t. the RS index increases; the sensitivity increases.

**Figure 9.**
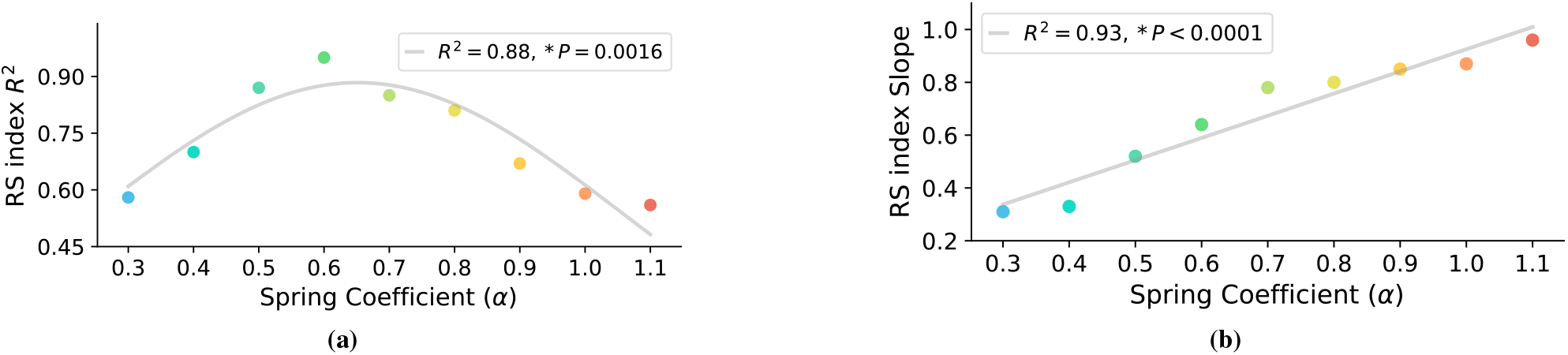
Correlation between R^2^ and slope of linear fitting for metabolic rate versus RS index. Figure (a) shows that the R^2^ of linear fitting for metabolic rate versus RS index is a quadratic diagram, and figure (b) indicates a linear relation between the slope of linear fitting for metabolic rate versus RS index in different spring coefficients. Based on figure (a), for the spring coefficients *α* < 0.3 and *α >* 1.1, R^2^ is very low; there is no linear relation for these ranges of *α*. Therefore, the RS index is only applicable for a range of spring coefficients (0.3 ≤ *α* ≤ 1.1) about the optimum spring coefficient (*α\** = 0.6). Figure (b) indicates that by increasing the spring coefficient, the sensitivity of the metabolic rate versus RS increases.

**Figure 10.**
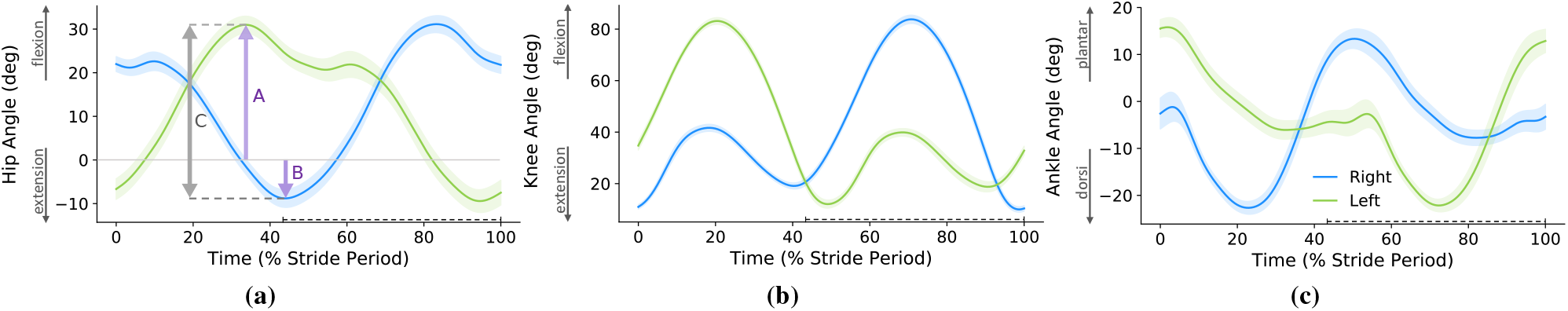
Joints’ trajectories and definition of kinematic indexes. In this figure, (a), (b), and (c) show the hip, knee, and ankle joints’ trajectories, respectively. The start of the gait cycle is at the heel-strike instant of the right leg, and the swing phase is shown with a dashed line over the horizontal axes. Figure (a) shows the hip joint trajectory and the definition of kinematic indexes. In figure (a), *A* is the maximum angle of the left hip joint, *B* is the minimum angle of the right hip joint, and *C* is the absolute value of the difference between *A* and *B*. It is important to note that *A* and *B* are vectors and can be either positive or negative, and *C* is a distance and it is always positive. As discussed in **A kinematic index for metabolic rate reduction** subsection, the kinematic indexes are defined as range of motion (RM) = *C* = *|A−B|*, symmetry of motion (SM) = *|A* + *B|^−1^*, and RS = RM × SM = *C ×|A* + *B|^−1^*; *|x|* means the absolute value of *x*.

RS is just a kinematic index and can be measured easily; in contrast to metabolic rate and EMG signals. It provides a metric not only for estimation of biological hip moment compensation and metabolic rate reduction but also for analyzing the quality of the subject’s training sessions. Using the proposed index, one can supervise the process of the training with the exoskeleton and lead the subjects to maximize their benefit from *I-RUN*. This index can also explain the individual differences in metabolic rate reduction of the experiments as well as an improvement over the sessions. Nevertheless, these findings are based on some simulations and extensive experiments are required for physical validations.

## Acknowledgements

The authors would like to appreciate the University of Tehran for providing support for this work. The Authors also would like to thank Mojtaba Rayati, Morteza Khosrotabar, and Elmira Aftabi for reading the first draft of this paper. Finally, our special gratitude goes to Saeid Amirimagham for providing the presented industrial design of *I-RUN* exoskeleton in this paper; Fig 1.

## Author contributions statement

H.A. conducted simulations and prepared figures. H.A., R.N., and M.N.A analyzed and interpreted the results. R.N. and H.A. drafted the manuscript. H.A., R.N., and M.N.A edited the manuscript. All authors reviewed the manuscript for important intellectual content.

## Footnotes

1 Hereafter by exoskeleton we mean an assistive exoskeleton devised for the metabolic rate reduction. We use exoskeleton and assistive device interchangeably.

2 The experimental markers are placed on anatomical landmarks of subjects.

3 The virtual markers are placed on the generic model.

4 The muscle activation pattern is the timing that muscles are activated or remain silent.

5 It is noteworthy to consider that RS is computed only for one stride. Obviously, if the subject performs a rhythmic running gait, these indexes are the same for whole strides. And, in the case that the subject performs a non-rhythmic running gait, the indexes of each stride should be studied individually; the overall indexes are the average indexes in each stride.

## References

1. Nasiri, R., Ahmadi, A. & Nili Ahmadabadi, M. Reducing the energy cost of human running using an unpowered exoskeleton. IEEE Transactions on Neural Syst. Rehabil. Eng. 26, 2026–2032 (2018).

2. Nasiri, R., Nili Ahmadabadi, M. & Ahmadi, A. Methods and systems for an exoskeleton to reduce a runners metabolic rate (U.S. Patent 10 549 138 B2, Feb. 2020).

3. Simpson, C. S. & et al. Connecting the legs with a spring improves human running economy. J. Exp. Biol. 222(2019).

4. Collins, S. H., Wiggin, M. B. & Sawicki, G. S. Reducing the energy cost of human walking using an unpowered exoskeleton. Nature 522, 212–215 (2015).

5. Van Dijk, W., Van der Kooij, H. & Hekman, E. A passive exoskeleton with artificial tendons: Design and experimental evaluation. In Rehabilitation Robotics (ICORR), 2011 IEEE International Conference on, 1–6 (IEEE, 2011).

6. Panizzolo, F. A. & et al. Reducing the energy cost of walking in older adults using a passive hip flexion device. J. Neuroeng. Rehabil. 16, 117 (2019).

7. Geyer, H. & Herr, H. A muscle-reflex model that encodes principles of legged mechanics produces human walking dynamics and muscle activities. IEEE Transactions on Neural Syst. Rehabil. Eng. 18, 263–273 (2010).

8. Song, S. & Geyer, H. Predictive neuromechanical simulations indicate why walking performance declines with ageing. J. Physiol. 596, 1199–1210 (2018).

9. Geijtenbeek, T. Scone: Open source software for predictive simulation of biological motion. J. Open Source Softw. 4, 1421 (2019).

10. Kutch, J. J. & Valero-Cuevas, F. J. Muscle redundancy does not imply robustness to muscle dysfunction. J. Biomech. 44, 1264–1270 (2011).

11. Karniel, A. & Inbar, G. F. Human motor control: learning to control a time-varying, nonlinear, many-to-one system. IEEE Transactions on Syst. Man, Cybern. Part C (Applications Rev. 30, 1–11 (2000).

12. Bernstein, N. A. The co-ordination and regulation of movements (Pergamon Press; Oxford, 1967).

13. Hug, T. & Tucker, K. Muscle coordination and the development of musculoskeletal disorders. Exerc. Sport Sci. Rev. 45, 1 (2019).

14. Horst, F. & et al. Explaining the unique nature of individual gait patterns with deep learning. Sci. Reports 9, 2391 (2019).

15. Koch, C. & Laurent, G. Complexity and the nervous system. Science 284, 96–98 (1999).

16. Power, J. D. & Petersen, S. E. Control-related systems in the human brain. Curr. Opin. Neurobiol. 23, 223–228 (2013).

17. Cestari, M., Sanz-Merodio, D., Arevalo, J. C. & Garcia, E. An adjustable compliant joint for lower-limb exoskeletons. IEEE/ASME Transactions on Mechatronics 20, 889–898 (2015).

18. Zhu, Y., Yang, J., Jin, H., Zang, X. & Zhao, J. Design and evaluation of a parallel-series elastic actuator for lower limb exoskeletons. In 2014 IEEE International Conference on Robotics and Automation (ICRA), 1335–1340 (2014).

19. Sawicki, G. S. & Khan, N. S. A simple model to estimate plantarflexor muscle–tendon mechanics and energetics during walking with elastic ankle exoskeletons. IEEE Transactions on Biomed. Eng. 63, 914–923 (2015).

20. Farris, D. J., Hicks, J. L., Delp, S. L. & Sawicki, G. S. Musculoskeletal modelling deconstructs the paradoxical effects of elastic ankle exoskeletons on plantar-flexor mechanics and energetics during hopping. J. Exp. Biol. 220, 4018–4028 (2014).

21. Uchida, T. K. et al. Simulating ideal assistive devices to reduce the metabolic cost of running. PLoS ONE 11, e0163417 (2016).

22. Dembia, C. L., Silder, A., Uchida, T. K., Hicks, J. L. & Delp, S. L. Simulating ideal assistive devices to reduce the metabolic cost of walking with heavy loads. PLoS ONE 12, e0180320 (2017).

23. Jackson, R. W., Dembia, C. L., Delp, S. L. & Collins, S. H. Muscle–tendon mechanics explain unexpected effects of exoskeleton assistance on metabolic rate during walking. J. Exp. Biol. 220, 2082–2095 (2017).

24. Lee, G. et al. Reducing the metabolic cost of running with a tethered soft exosuit. Sci. Robot. 2, eaan6708 (2017).

25. Nasiri, R., Rayati, M. & Nili Ahmadabadi, M. Feedback from mono-articular muscles is sufficient for exoskeleton torque adaptation. IEEE Transactions on Neural Syst. Rehabil. Eng. 27, 2097–2106 (2019).

26. Hamner, S. R. & Delp, S. L. Muscle contributions to fore-aft and vertical body mass center accelerations over a range of running speeds. J. Biomech. 46, 780–787 (2013).

27. Delp, S. L. & et al. Opensim: open-source software to create and analyze dynamic simulations of movement. IEEE Transactions on Biomed. Eng. 54, 1940–1950 (2007).

28. Schache, A. G. & et al. Lower-limb muscular strategies for increasing running speeds. J. Orthop. Sports Phys. Ther. 44, 813–824 (2014).

29. Hamner, S. R., Seth, A. & Delp, S. L. Muscle contributions to propulsion and support during running. J. Biomech. 43, 2709–2716 (2010).

30. Thelen, D. G. Adjustment of muscle mechanics model parameters to simulate dynamic contractions in older adults. J. Biomech. 125, 70–77 (2003).

31. Thelen, D. G. & Anderson, F. C. Using computed muscle control to generate forward dynamic simulations of human walking from experimental data. J. Biomech. 39, 1107–1115 (2006).

32. Hicks, J. Computed Muscle Control Theory (2018). Available at: https://stanford.io/2IKFMXT (Jun. 2018).

33. Uchida, T. K., Hicks, J. L., Dembia, C. L. & Delp, S. L. Stretching your energetic budget: how tendon compliance affects the metabolic cost of running. PLoS ONE 11, e0150378 (2016).

34. Crowninshield, R. Use of optimization techniques to predict muscle forces. J. Biomech. Eng. 100, 88–92 (1978).

35. Pedotti, A., Krishnan, V. & Stark, L. Optimization of muscle-force sequencing in human locomotion. J. Biomech. Eng. 38, 57–76 (1978).

36. Bregman, D. J. & et al. The effect of ankle foot orthosis stiffness on the energy cost of walking: a simulation study. Clin. Biomech. 26, 955–961 (2011).

37. Hicks, J. L. & et al. Is my model good enough? best practices for verification and validation of musculoskeletal models and simulations of movement. J. Biomech. Eng 137,: 020905 (2015).

38. Hicks, J. OpenSim User’s Guide CMC Best Practices (2012). Available at: https://stanford.io/38YtN3t (Jan. 2012).

39. Roberts, T. J. & Gabaldón, A. M. Interpreting muscle function from emg: lessons learned from direct measurements of muscle force. Integr. Comp. Biol. 48, 312–320 (2008).

40. De Luca, C. J. The use of surface electromyography in biomechanics. J. Appl. Biomech. 13, 135–163 (1997).

41. Ding, Y., Kim, M., Kuindersma, S. & Walsh, C. J. Human-in-the-loop optimization of hip assistance with a soft exosuit during walking. Sci. Robotics 3, eaar5438 (2018).

42. Zhang, J. et al. Human-in-the-loop optimization of exoskeleton assistance during walking. Science 356, 1280–1284 (2017).

